# Soil to human health continuum: Exploring ergothioneine and mycorrhizal fungi in shaping the wheat microbiome

**DOI:** 10.1101/2025.05.28.656623

**Authors:** Sri Sai Nandini Ravi, Alexandra Pipinos, Nick Insley, Jinjun Kan, Wade Heller, Andrew Smith, Gladis Zinati, John Richie, Harsh Bais

## Abstract

**Background:** The association between plants and soil microbes is critical for both soil and plant health. Studies have shown that introducing beneficial microbial inoculants can shape the soil microbiome community for plant health. Among these microbes, mycorrhizal fungi play a well-documented role in enhancing nutrient uptake in plants. Ergothioneine (ERGO), a compound well-known for its anti-inflammatory and antioxidant properties, has been linked to increased longevity in various model systems and its significance for human health. However, neither animals nor plants contain ERGO biosynthetic pathways, which are limited to fungi, including and some species of bacteria, including Actinomyceota, Cyanobacteria, and Methylobacteria. Though the leading dietary sources of ERGO for humans are fungi in the form of mushrooms or fermented foods, biofortification of crops by promoting the production and uptake of ERGO from microbial sources in the soil has promise for enhancing nutritional quality and public health outcomes.

**Results:** This study explores the of interaction between soil ERGO application and arbuscular mycorrhizal fungi (AMF) in plant-symbiotic relationships to increase the ERGO content in the staple crop wheat (*Triticum aestivum*). We investigate how ERGO supplementation, both alone and in combination with AMF, influences the wheat root and soil microbiome in a greenhouse experiments. Our data shows that plants can take up ERGO in absence of AMF fungi. In addition, treatment with pure ERGO and ERGO in combination with AMF altered microbial diversity and community structure in both the rhizosphere and rhizoplane regions of wheat roots.

**Conclusions:** Overall, our work reveals that plants can readily take up ERGO from soil, both with and without AMF presence, highlighting a broader role of ERGO in connecting soil health to human health, a connection that warrants further investigation.

## Background

In recent years, a decline in crude protein content in agricultural crops has been triggered by several factors, including plant breeding, climate change, decline in soil organic matter, and changes in environmental legislation [1,2]. Literature has stated that approximately 2.4-2.9% of crude protein content in grain crops has been lost over the past two decades [1]. Meta-analyses have also found that rising environmental CO_2_ levels contribute to decreased protein content in staple crops by restricting nitrogen uptake and inhibiting protein biosynthesis [3]. Furthermore, the protein quality in plants is generally lower due to its lesser quantities of essential amino acids, specifically lysine and sulfur-containing amino acids [4]/ It is widely known that the net protein content in plants is influenced by the overall availability of Nitrogen (N) in the soil. sources rhizobia Recent studies increasingly focus on addressing the lack of protein content by exploring the concept of holistic microbiome approaches in plants [5]. Studies have shown that plant-associated microbiomes may contribute significantly to nutrient assimilation, including N in non-leguminous crops. However, it remains unclear what specific microbial population is influenced by inoculation with arbuscular mycorrhizal fungi (AMF), ergothioneine (ERGO), or their combination, and which specific taxa contribute to nitrogen assimilation and how nitrogen assimilation and protein formation are influenced by microbial interactions in the soil. Terrestrial plants associate closely with arbuscular mycorrhizal fungi (AMF), which enhance root surface area, while also facilitating phosphorus (P) uptake [6]. Additionally, AMF have been shown to influence the uptake of organic nitrogen compounds, including amino acids [7]; however, the definite role of mycorrhizal associations in enhancing protein content in plants is not well established. Recent studies have demonstrated that fungal species, including mushrooms, can biosynthesize an unusual amino acid called ergothioneine (hereafter ERGO), which has been recognized for its nutraceutical properties in various model systems, and serves as an example of a bioactive compound that may be influenced by beneficial microorganisms in the rhizosphere, including AMF [8]

ERGO was first discovered in 1909 by Charles Tanret, who extracted the amino acid from the rye grain infected by ergot fungus, *Claviceps purpurea* [9]. In recent literature, ERGO has been defined as a “longevity vitamin” due to its anti-inflammatory and antioxidant impacts pertaining to human health [10]. ERGO has been linked to several studies highlighting its relevance as an antioxidant with cytoprotective properties [11]. ERGO commonly accumulates in tissues vulnerable to oxidative stresses, making it a key focus in human health research, particularly in relation to cardiovascular disease, liver disease, and neurodegenerative diseases [12]. However, ERGO is present as a naturally occurring antioxidant in plants and mammals [9], yet it has been found that neither animals nor plants are capable of biosynthesizing it [13].

Fungi utilize a two-gene pathway, *Egt1* and *Egt2* [14,15], to produce ERGO from the precursor amino acid histidine, while bacterial biosynthesis requires a five-gene pathway *EgtA-E* [16]. The known prokaryotes that biosynthesize ERGO are Actinobacteria, cyanobacteria, and methylobacteria species [17]. Fungal speciesincluding *Neurospora crassa, Cordyceps militaris*, and mushroom fruiting bodies also produce ERGO. Previous studies have detected various quantities of the amino acid in the small grain crops oats, wheat, and barley [18], while other studies measuring ERGO in small grains found that root inoculation with specific AMF species had higher ERGO levels than a non-inoculated control. Thus, there is the potential for a connection between plant accumulation or uptake of ERGO and the presence of fungal and/or bacterial species in the soil [18] Supporting this, recent studies have shown that crops grown with AMF exhibit enhanced ERGO content across multiple plant species [8]. Although a specific transporter for ERGO in animals has been [19] the exact transport and uptake mechanism of ERGO in plants remains unknown. Therefore, further research is needed to elucidate how ERGO is taken up in cereal crops to enhance its levels for increased dietary intake in humans. Simultaneously, there is limited understanding of how soil and plant microbiomes respond and adapt to ERGO supplementation and AMF inoculation, warranting further research.

To further our understanding of how the root and soil microbiome influence ERGO uptake in plants, we conducted a seed-to-seed experiment using the following soil treatments: AMF inoculation, ERGO supplementation, combined AMF/ERGO treatment, and untreated control plants. We hypothesized that association with AMF will stimulate ERGO uptake in plants. Additionally, the presence of both ERGO and AMF in the soil may further enhance ERGO uptake. We also predicted that AMF association, with or without ERGO, could influence the plant-associated and soil microbiome and enrich microbial taxa potentially involved in ERGO uptake. To assess the changes in microbiome structure in response to the treatments, amplicon high throughput sequencing was performed for the bulk and rhizosphere soil from all treatment groups. Interestingly, our data showed that plants were able to take up ERGO with/without AMF, suggesting the presence of an efficient uptake system in plants for ERGO exists. In addition, analysis of microbiome diversity and community structure in the rhizoplane and rhizosphere revealed that ERGO supplementation-with or without AMF drove microbial enrichment in the wheat plant rhizosphere.

## Methods

### Chemicals

L-ergothioneine (L-ERGO, purity > 99.99+%) was obtained from MedChemExpress^®^ (Monmouth Junction, NJ). L-(+)-ergothioneine (ERGO) standard was purchased from Toronto Research Chemicals (Toronto, ON, CA), and L-(+)-ergothioneine-d9 (ERGO-d9) was purchased from Santa Cruz Biotechnology (Dallas, TX, USA) as internal standard. The ERGO inoculum for dose-response experiments was prepared by dissolving L-ERGO into 30 mL of nanopore water and further diluting the solution into 0.125 mg/mL and 0.375 mg/mL concentrations. These two exogenous concentrations were used based on the natural titers of ERGO found in biological or soil systems.

### Arbuscular mycorrhizal fungi

A mixed species consortium of AMF. These include consortium of AMF *Claroideoglomus etunicatum, Claroideoglomus claroideum, Rhizophagus irregularis, Rhizophagus intraradices*, and *Funneliformis mosseae* was propagated from agricultural soil under organic management at the Rodale Institute, Kutztown, PA. To prepare the inoculum, field soil was diluted 1:10 in a mixture of autoclaved sand, soil, vermiculite, and turface (Applied Industrial Materials, Corp., Deerfield, IL, 60015) (1:0.75:1:0.75 v/v), filled into 66-ml containers into which individual seedlings of Bahia grass (*Paspalum notatum*) were transplanted. After 3 months growth in the greenhouse, the plants and soil were transferred into 7-gallon grow-bags filled with a 3:1 mixture of plant-based compost and grown outdoors on raised benches over the summer and autumn to allow the spores to mature as in (Douds et al. (2010). Prior to utilization in the experiment, roots were cut into 1-cm lengths and a spore density was calculated to be approximately 100 spores/ml. “Mock inoculum”, to control for the additional nutrients contained with the compost, was prepared by autoclaving inoculum at 121°C for 60 minutes.

### Plant materials and growth conditions

Spring wheat (*Triticum aestivum*; v) cv. Surpass seeds were Johnny’s Selected Seeds (Winslow, ME) used for all experiments. The seeds were directly sown into 1-L 3.81 cm deep into 8.89 x 8.89 x 12.7 cm pots containing pre-moistened potting media (ProMix, City, State) without mycorrhizae and topped with a thin layer of vermiculite. The plants were grown in a greenhouse at 23°C with a 16-hour photoperiod, and a Photosynthetic Photon Flux (PPF) of 2000 µmol/s.

### ERGO and AMF treatment of *Triticum aestivum* and ergothioneine in microbiome analysis

This experiment aimed to evaluate plant growth and soil microbiome responses to various treatments involving AMF and ERGO, with results compared across different growth phases of spring wheat. Germinating spring wheat seedlings were replanted into 160 pots containing ProMix without mycorrhizae. The following treatments were evenly divided amongst the wheat plants: +mixed AMF species/- ERGO, +mixed AMF species/+ ERGO, -mixed AMF species/+ERGO, and –mixed AMF species/-ERGO (control). For plants receiving the AMF treatments, 60 ml60 ml of inoculum were inoculated onto the spring wheat seedlings, while plants that did not receive mixed AMF species received a containing ∼6,000 spores was homogenized with 40-L ProMix BX (PremierTech Horticulture, Riviere-du-Loup, Quebec, CA) and used to fill each set of 40 pots/treatment (∼150 spores/pot), and mock inoculum that consisted of autoclaved inoculum added at the same rate to pots for the non-AMF treatments. For the ERGO dosing, ERGO treatments were dosed with 0.025mg L-ergothioneine/pot, while control plants were dosed with mock inoculum that consisted of distilled water only. All 160 pots were then topped with a thin layer of vermiculite and thoroughly watered. that ERGO was applied on day 0 and boosted with 0.025 mg/pot ERGO on day 40 of growth. Thus, for days 40, 50, and 60 harvest, 13 plants per treatment were harvested, for a total of 52 plants per day of harvest, and an overall total of 152 plants for all 4 treatments and 3 harvest days. For plant growth, images were taken, and plant samples were collected on for the day 60 harvest, where plants were divided between roots and shoot and placed in separate bags. For microbiome analysis, the roots were gently shaken for rhizosphere soil collection from the soil surrounding/attached to them. Bulk soil was collected from pots where plant roots did not expand. The plant roots, rhizosphere soil and bulk soil samples were sent to Stroud Water Research Center (Avondale, PA), for microbial community analyses (described separately below). For ERGO analysis, the roots, shoots, and grain of each plant were harvested and placed in an oven dryer at 60 ℃ for at least 24 hours. For ERGO quantification, the dried samples were ground to a powder using a small sample grinder coupled with a size 40 mesh (0.4 mm). The finely ground samples were then sent to Penn State College of Medicine for analysis by High-Performance Liquid Chromatography (HPLC) as described in (REF).

### ERGO Analysis and Sample Preparation Procedure

ERGO and ERGO-d9 stock solutions (1 mg/mL) were prepared in methanol. The standard working solutions were prepared with a series of dilutions of the standard stock solution in methanol/water (50/50), resulting in concentrations from 10 to 10,000 ng/mL. ERGO-d9 working solution was also prepared by diluting its stock solution using methanol/water (50/50). The ERGO standard curve was prepared by spiking an internal standard working solution into 50% methanol to make the standard concentrations from 1 to 1000 ng/mL. The standard curves were constructed by plotting the ratio of the peak area of ERGO.

The grain, root, and shoot samples were ground into powder before processing. After weighing the powders (around 25 mg), 10 μL of the internal standard working solution was spiked, and 990 μL of 50 % methanol was used to extract ERGO from the powder. The mixture was vortexed for 20 min under room temperature, then centrifuged at 4 °C, 8765 x *g* for 10 min. After transferring the supernatant to another vial, the pellet was re-extracted by 1 mL of 50% methanol. An equal volume of each supernatant was combined and vortexed before being loaded onto an HPLC-MS/MS system for analysis.

### Ultra-Performance Liquid Chromatography HPLC-MS/MS Analysis

ERGO was analyzed using a Sciex 6500+ QTrap mass spectrometer coupled with an ExionLC HPLC separation system. A 1.7 μm ACQUITY UPLC BEH C18 analytical column (2.1 × 100 mm, Waters, Ireland) was used with the following mobile phase: mobile phase A consisted of 0.1% formic acid in water, and mobile phase B consisted of acetonitrile. A gradient program was applied to separate ERGO from the matrix and other impurities. The Sciex 6500+ QTrap mass spectrometer was equipped with an electrospray ionization probe in positive mode. The ion spray voltage was 5500 V, the temperature was 500 °C, the ion source gas 1 was 20 L/h, and the ion source gas 2 was 20 L/h. The multiple reaction monitoring mode was used to analyze and quantify ERGO and ERGO-d9, with transitions of m/z at 230 > 127 for ERGO and m/z at 239 > 127 for ERGO-d9. All peaks were integrated and quantified by Sciex OS 1.5.

### DNA extraction and Sequencing

Genomic DNA fro roots, rhizosphere and bulk soil samples were extracted by using DNeasy PowerSoil Pro kits (Qiagen, Hilden, Germany) following the manufacturer’s instructions. DNA quantity and quality were measured using an ND-2000 Nanodrop spectrometer (Thermo Fisher Scientific, Waltham, USA). Amplicon high throughput sequencing was conducted to characterize detailed bacteria/archaea and fungi communities. For bacteria and archaea, the V4 variable region of the 16S rRNA genes was amplified using 515f (5′-GTGYCAGCMGCCGCGGTAA-3′; [20] and 806r (5′-GGACTA CNVGGGTWTCTAAT-3′; (21) following the Earth Microbiome Project protocol [22]. For fungi, the ITS2 region was amplified using ITS3-F (5’-GCATCGATGAAGAACGCAGC-3′) and ITS4-R (5’-TCCTCCGCTTATTGATATGC-3′[23]. 16S rRNA genes were amplified with following thermocycling program: 5 min at 94°C for initialization; 30 cycles of 30 s denaturation at 94°C, 30 s annealing at 53°C, and 30 s extension at 72°C; followed by 8 min final elongation at 72°C. Fungal ITS regions were amplified with 3 min at 95°C for initialization; 33 cycles of 20 s denaturation at 95°C, 20 s annealing at 56°C, and 30 s extension at 72°C; followed by 5 min final elongation at 72°C. Sequencing libraries were prepared by using NEBNext Ultra II DNA Library Prep Kit for Illumina (New England Biolabs, Massachusetts, United States) following manufacturer’s recommendations. High-throughput sequencing was performed at Magigene (Magigene Biotechnology, Guangzhou, China) on an Illumina Nova6000 platform (paired-end 250-bp mode). Raw sequencing data obtained in this study are deposited to GenBank database under the accession number xxxx

### Statistical Analysis of Plant Traits

For statistical analyses of plant traits, a two-way ANOVA test was run to compare each treatment group with the control group. Dunnett multiple comparisons test was conducted for hypothesis testing. To analyze ERGO accumulation in root, shoot, and grain, a two-way ANOVA test was run to compare treatments groups to control samples amongst plant tissue type. Dunnett multiple comparisons test was also conducted for hypothesis testing. All tests show significant results at p < *a* = 0.05.

### Microbiome Diversity and community analysis

Raw Illumina sequences were processed with the QIIME 2 software package (version 2021.11) [24]. After demultiplexing, all raw sequence reads were carried out with quality control, denoising, filtering, merging, and chimera removal through q2-DADA2. Amplicon sequence variants (ASVs) were generated, and a Naïve Bayes classifier artifact2 was applied to assign the ASVs to taxa at 99% using the Silva classifier 132 [25] for 16S rRNA genes and UNITE version 8.2 (February 20, 2020) for ITS regions. The ASVs were normalized by rarefaction approach performed with Qiime 2 pipeline, with cutoffs at 52,000 sequences for bacteria/ archaea and 6,400 for fungi. Both alpha and beta diversity metrics were analyzed to assess microbial diversity across various treatments and sample types (root, rhizosphere, and soil). Alpha diversity was evaluated using Shannon diversity for each treatment within the sample types. A one-way analysis of variance (ANOVA) was performed on Shannon diversity indices, and Post hoc comparisons were performed using Tukey’s Honest Significance Difference (Tukey HSD) test [26,27], to identify significant pairwise differences using plyloseq, ggplot2, dplyr, multcompView, tidyr, and viridis in RStudio. Beta diversity was examined through Bray-Curtis dissimilarity matrices using the ordinate function in phyloseq[28] derived from ASV abundance data to investigate variations in microbiome composition. Principal Coordinates Analysis (PCoA) was conducted on the Bray-Curtis matrix to visualize the segregation of microbial communities based on treatment and sample type. The statistical significance of differences in community composition was tested using permutational multivariate analysis of variance (PERMANOVA) with 999 permutations, using the adonis2() function from the vegan package, focusing on the effects of treatments and sample types on microbiome structure. Additionally, PERMANOVA was employed to identify significant changes between individual treatment pairs and sample type [29]

### Indicator Species analysis

Indicator Species analysis was performed using the multipatt() function from indicator species package in R with 16s and ITS ASVs to identify microbial taxa significantly abundant at different sample types based on the applied treatments [30]. Only species with p <0.05 were retained. Relative abundance was calculated by normalizing ASV count per sample and average treatment groups. Bubble plots and summary tables are generated for 16s and ITS data [31]. Each bubble plot represented a significant species-treatment associations, where the size of the bubble plot reflected the mean relative abundance, and the fill color denoted the significance (p-values). Indicator values were displayed to each species for added interpretability. In addition to that, a summary table of indicator species analysis for both 16s and ITS were generated with its treatment association, relative abundance and p-value. This format provides a concise overview of taxa that were statistically significant and biologically meaningful

### Correlation network Analysis

Correlation network analysis was conducted to explore correlation between added AMF species under M and ME treatments and their influence on bacterial abundance. Out of 5 AMF mixed species only 3 were found to be present in ASVs. Spearman Correlation coefficients were calculated between the present AMF species in ASVs and other bacterial taxa using Hmisc package in R[32]. Correlation with p-value < 0.05 were retained. Separate correlation matrices were generated for each treatment and domain (16S and ITS). Significant positive and negative correlations were extracted and formatted into edges and nodes lists. These lists were imported to Gephi for visualization [33]. The Furchterman-Reingold Algorithm was used to optimize mode positioning for network clarity [34]

## Results

### Treatment of ergothioneine and mycorrhizae influence plant biomass in *Triticum aestivum*

Wheat plants were harvested on day 60 and were analyzed for biomass for all treatment groups including controls (See Figure 1). The data for the biomass accumulation in wheat plants treated with pure. Inoculation with ERGO, mycorrhizae and the combination of ERGO and mycorrhizae (ME) showed increase over the untreated plants (Figure 1 and SUPPLEMENTARY ONLINE Figure 1). Based on the results, the plants treated with ‘ERGO’ contained the highest average plant weight increased plant biomass compared to an untreated control (Figure 1). ERGO and mycorrhizae alone resulted in higher biomass accumulation than the combination of ERGO and Mycorrhizae (ME). The data for morphometric plants showed that both ERGO (E) and mycorrhizae (M) may play a critical role in enhancing biomass in wheat plants.

**Figure 1:**
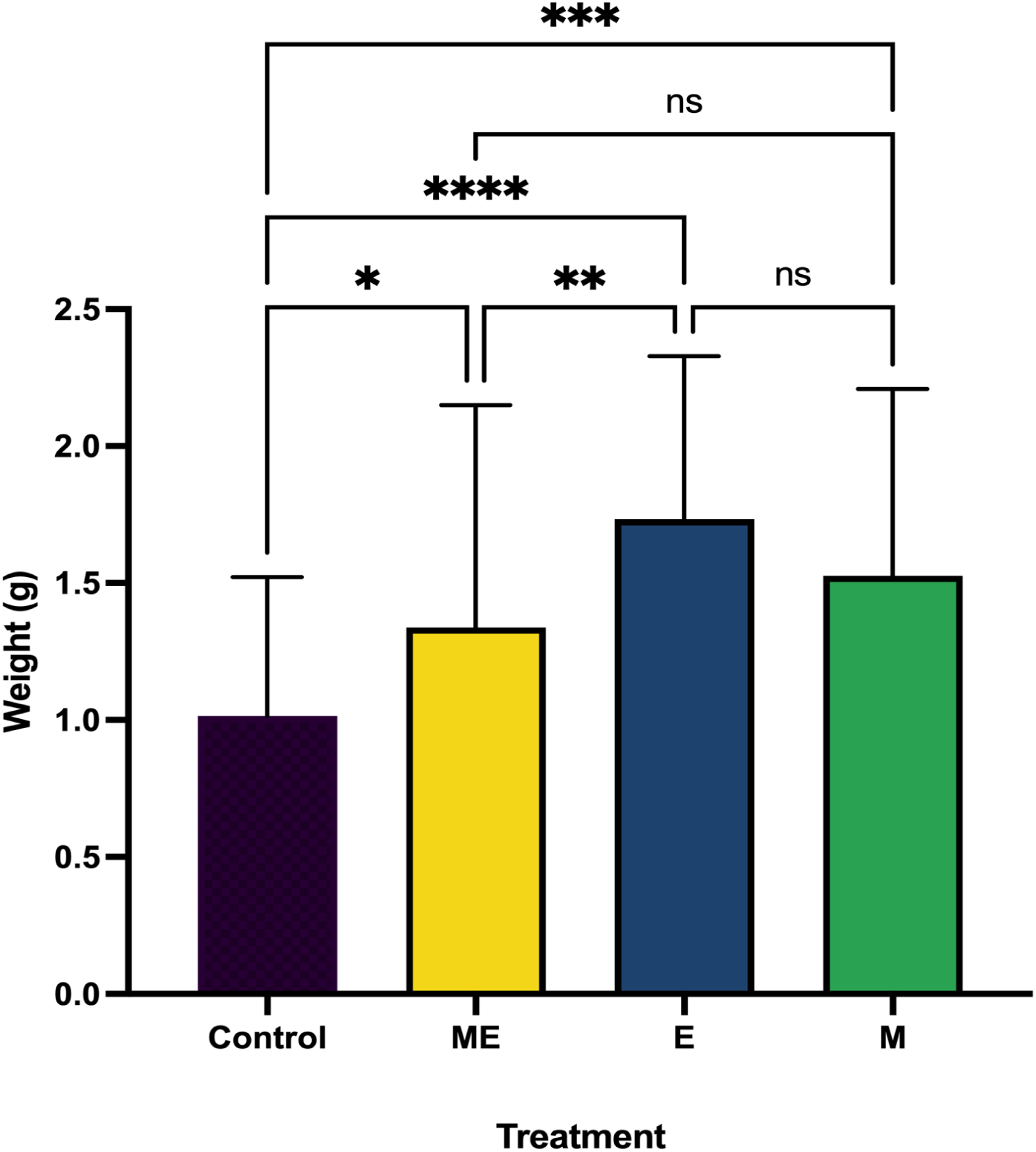
Biomass in spring wheat plants inoculated with ergothioneine (E), mycorrhizal fungi (M), and both (ME). The overall biomass was calculated on day 60 post-harvest. Control samples are not treated with either mycorrhizal species or ergothioneine. Asterisks indicate statistically significant differences (*p < 0.05, **p < 0.01, ***p < 0.001, ****p < 0.0001) between treatments.

### Ergothioneine accumulation in plants treated with ERGO and mycorrhizal inoculations

The ERGO levels in *Triticum aestivum* root, shoot, and grain were displayed in Figure 2. Analysis revealed a significant increase in the ERGO accumulation in roots under the pure exogenously amended ERGO’ treatment, with mycorrhizae and ERGO in the ‘ME’ treatment resulted in a significant increase in ERGO uptake in both roots and shoots compared to control samples (See Figure 2). In the grains, although a slight increase in ERGO concentration levels was observed compared to controls, this difference was not statistically significant. In *T. aestivum* plants treated with ERGO (E), the highest and most significant ERGO content was observed in the roots compared to other treatments, with a similar trend also noted in the shoots. Interestingly, the amount of ERGO detected in the grains of ‘E’-treated plants was lower than that observed in the control group. Lastly, in plants treated with mycorrhizae alone (M), ERGO levels were lower than those in the control group across roots, shoots, and grains. Our data shows that plants treated with either ERGO or with a co-inoculation of ERGO and mycorrhizae showed temporal uptake of ERGO in shoots and roots.

**Figure 2:**
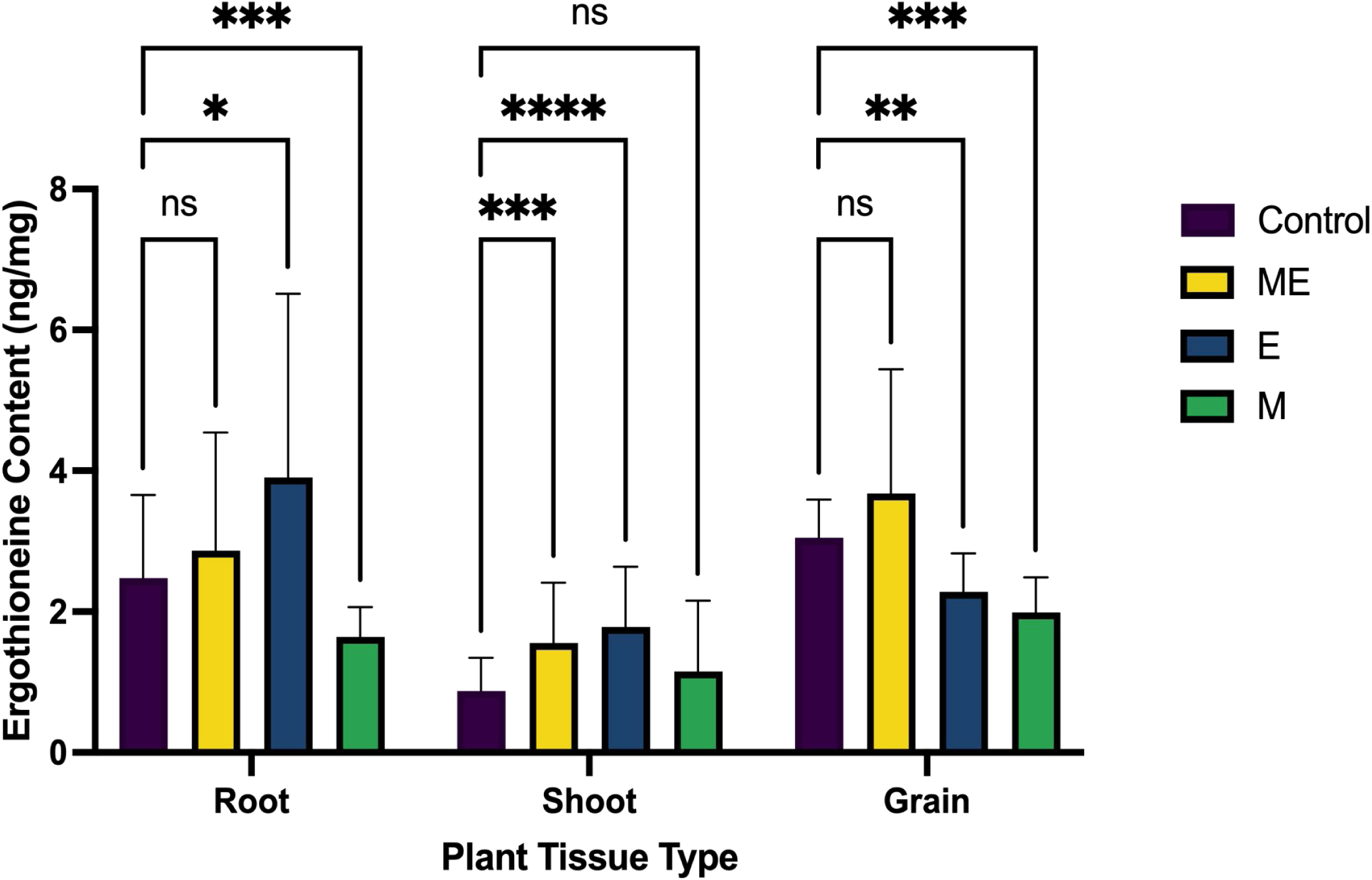
Ergothioneine content in wheat plants inoculated with ergothioneine (E), mycorrhizal fungi (M), and both (ME). Plants were harvested in day 60 and ergothioneine content was analyzed in roots, shoots and grains. Asterisks indicate statistically significant differences (*p < 0.05, **p < 0.01, ***p < 0.001, ****p < 0.0001) between treatments.

### Impact of ERGO and co-inoculation of ERGO and mycorrhizal fungi on soil microbiome assembly

The Shannon diversity plots for the bacteria/archaea and fungi showed differences across the treatments and the from different sampling zones differed between treatments (Figure 3, 4). With a p-value < 0.05, annotations of significance above each boxplots showed the differences in microbial diversity within the supplementation and mycorrhizal (M) treatment may modulate the rhizosphere microbiome assembly in the wheat plants. Contrary to the bacterial diversity index for the treatments across the sampling zones, the overall fungal diversity was not significantly different across all the sample zones (root plane, rhizosphere, and bulk soil) across all the treatments (Figure 4).

**Figure 3:**
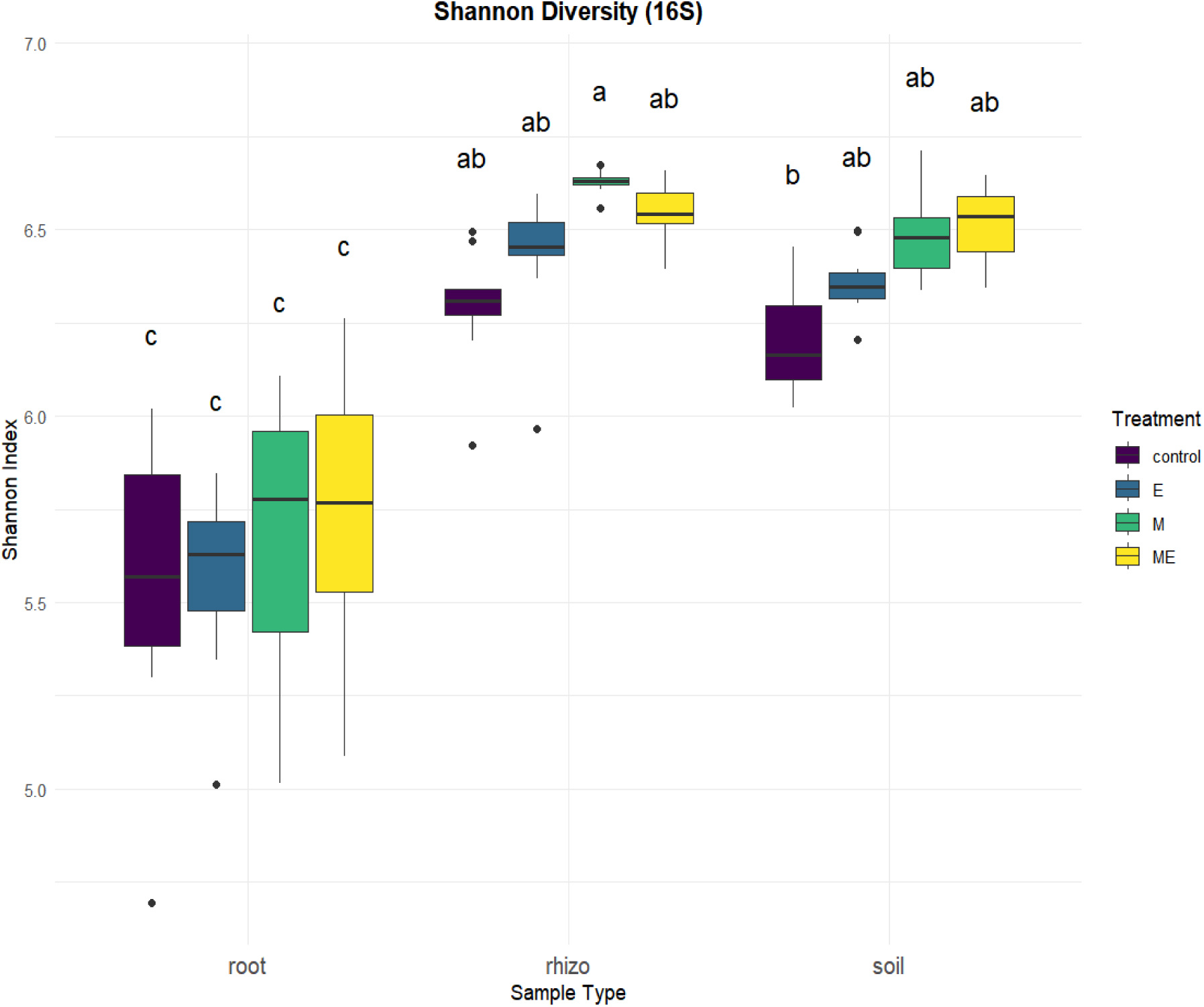
Boxplot illustrating the Shannon diversity index of bacteria (16S ASVs) across different treatments and sample types (root, rhizosphere, and bulk soil). The data reveals variations in bacterial diversity levels based on treatment conditions, with distinct distributions evident for each sample type with a p-value < 0.05. The letters above denotes the statistical significance among treatment groups within each sample type, based on post-hoc Tukey HSD tests. Control samples are not treated with either mycorrhizal species or ergothioneine. The other treatment depicts: ‘ME’ treatment in represents wheat plants inoculated with both mycorrhizal species and ergothioneine. ‘E’ treatment represents wheat plants inoculated with ergothioneine. ‘M’ treatment in represents wheat plants inoculated with mycorrhizal species.

**Figure 4:**
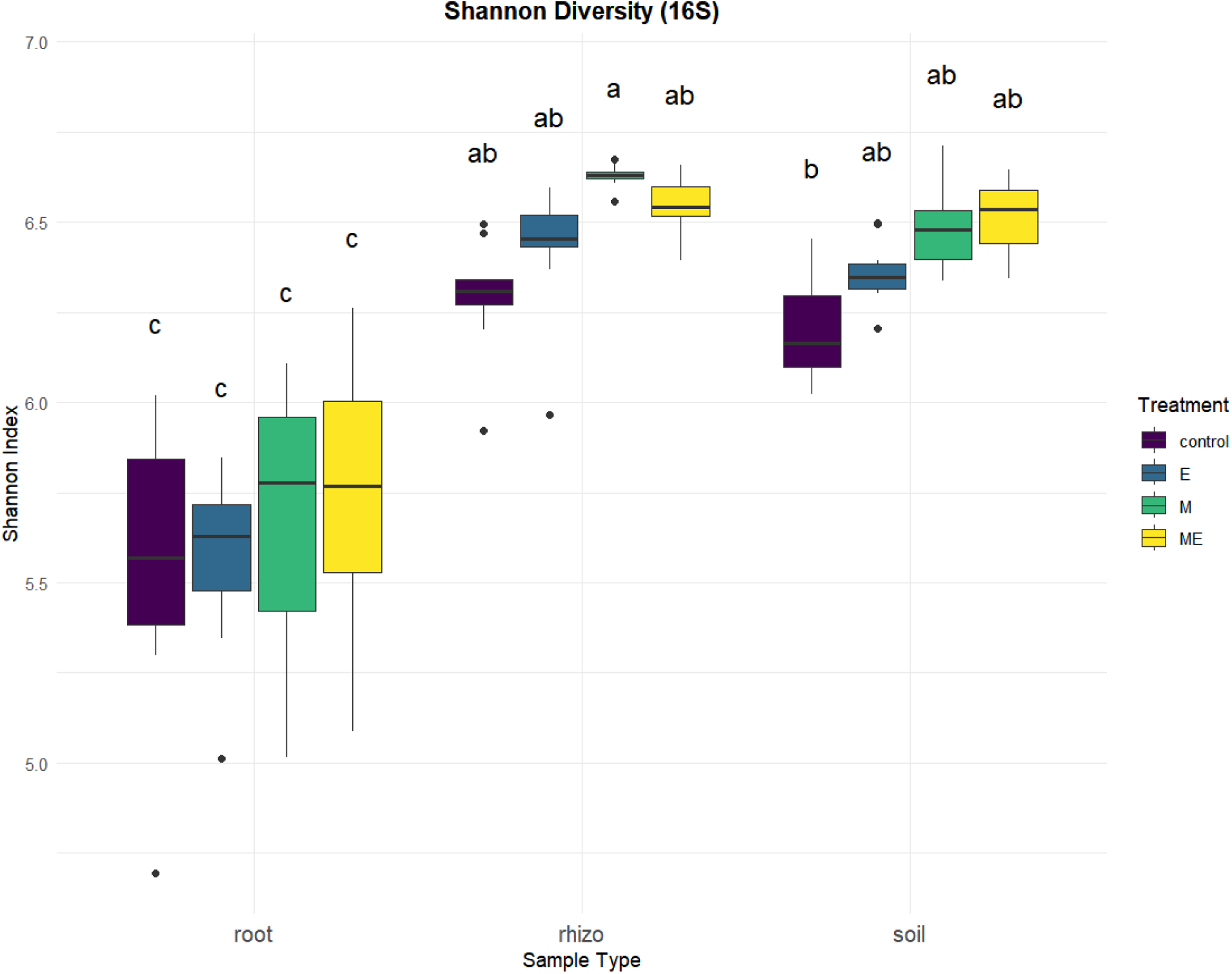
Boxplot illustrating the Shannon diversity index of bacteria (ITS ASVs) across different treatments and sample types (root, rhizosphere, and bulk soil). The data reveals variations in bacterial diversity levels based on treatment conditions, with distinct distributions evident for each sample type with a p-value < 0.05. The letters above denotes the statistical significance among treatment groups within each sample type, based on post-hoc Tukey HSD tests. Control samples are not treated with either mycorrhizal species or ergothioneine. The other treatment depicts: ‘ME’ treatment in represents wheat plants inoculated with both mycorrhizal species and ergothioneine. ‘E’ treatment represents wheat plants inoculated with ergothioneine. ‘M’ treatment in represents wheat plants inoculated with mycorrhizal species.

The impact of exogenous ERGO and mycorrhizae or both at different sample zones on assembly of bacterial and fungal communities (β-diversity) across different sampling zones (root plane, rhizosphere, and bulk soil) were shown in Figure 5-6. Permutational Multivariate Analysis of Variance (PERMANOVA) analysis indicated that microbiome was significantly influenced by ERGO, mycorrhizae and ERGO (ME) co-inoculation (p = 0.001 for 16S and p = 0.002 for ITS) (Figure 5, 6). Treatment accounted for approximately 21.4% of the observed differences, while sample type (root, rhizosphere, and soil) explained 18.1% of bacterial data. Whereas in fungal data, around 7.3% of the observed differences were due to treatment and 20% difference in abundance levels was related to type of sample.

**Figure 5:**
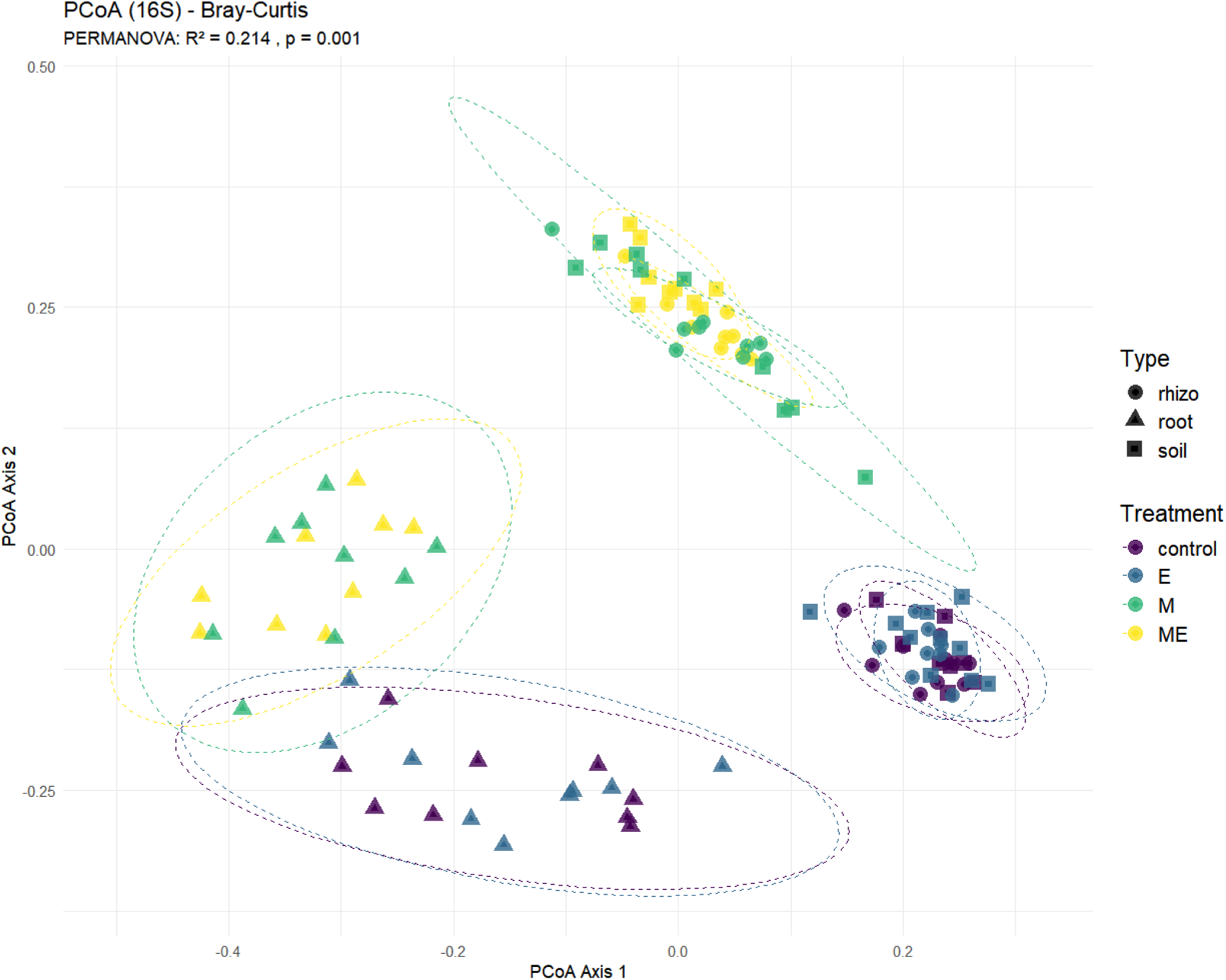
Bray-Curtis dissimilarity index and PCoA (Principal Coordinates Analysis) analysis of the 16S ASV data shows the relationships between the bacterial communities from several sample types (bulk soil, rhizosphere, and root plane) across different treatments with a p-value of 0.001. Every point on the plot represents a sample; various colors denote specific treatments, and various shapes indicate different sorts of samples. The spots’ spatial organization shows clustering patterns, indicating that sample type and treatment impact the composition of the microbial community. Control samples are not treated with either mycorrhizal species or ergothioneine. The other treatment depicts: ‘+M+E’ treatment in represents wheat plants inoculated with both mycorrhizal species and ergothioneine. ‘E’ treatment represents wheat plants inoculated with ergothioneine. ‘M’ treatment in represents wheat plants inoculated with mycorrhizal species.

**Figure 6:**
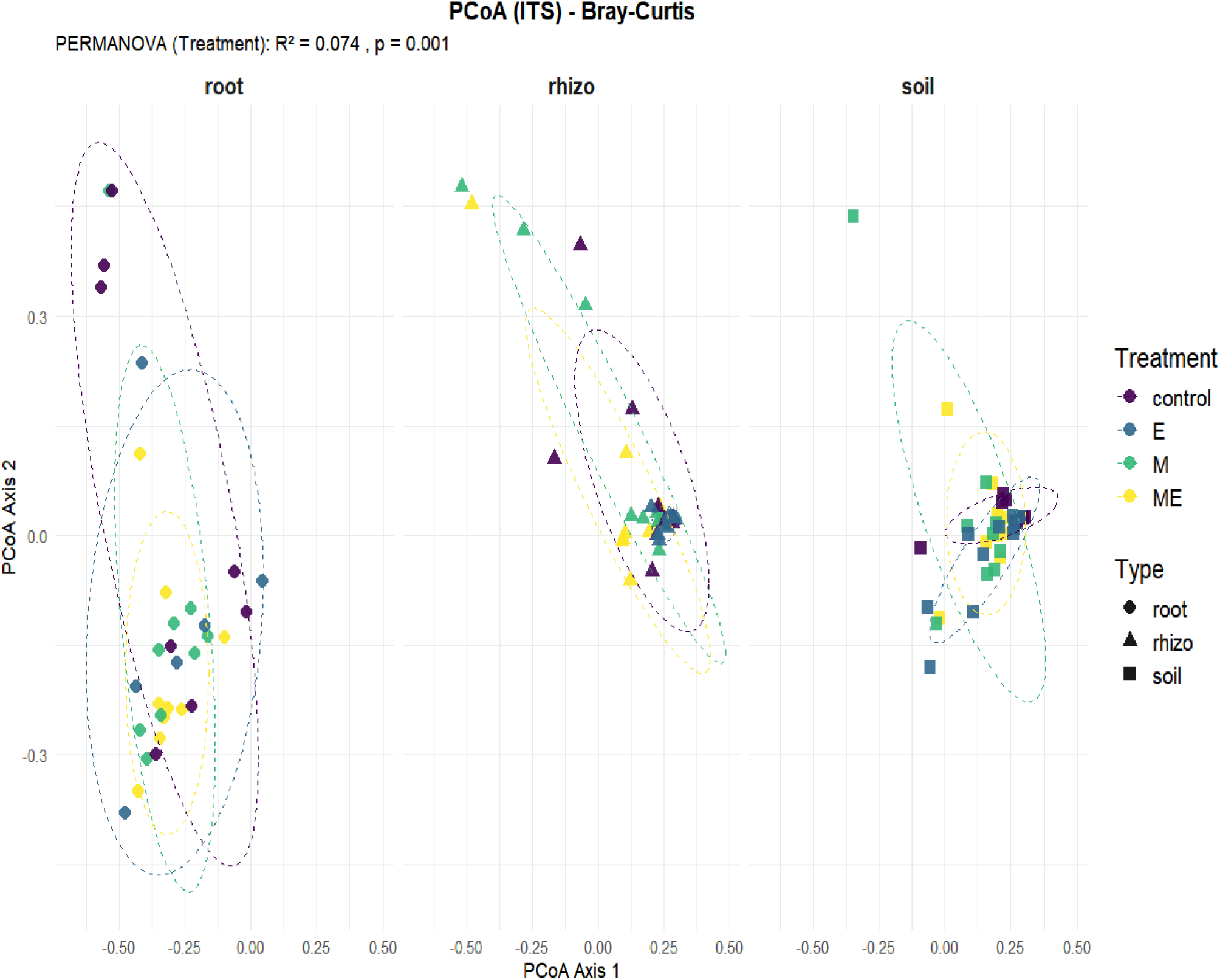
Bray-Curtis dissimilarity index, the PCoA (Principal Coordinates Analysis) analysis of the ITS ASV data shows the relationships between the fungal communities from several sample types (soil, rhizosphere, and root plane) across different treatments with a p-value of 0.002. Every point on the plot represents a sample; various colors denote treatments, and various shapes indicate different sorts of samples. The spots’ spatial organization shows clustering patterns, indicating that sample type and treatment impact the composition of the microbial community. Control samples are not treated with either mycorrhizal species or ergothioneine. The other treatment depicts: ‘+M+E’ treatment in represents wheat plants inoculated with both mycorrhizal species and ergothioneine. ‘E’ treatment represents wheat plants inoculated with ergothioneine. ‘M’ treatment in represents wheat plants inoculated with mycorrhizal species.

Significant changes in the composition of the bacterial communities were observed when ERGO and mycorrhizae (ME) were added to the wheat rhizosphere compared to the untreated control (Figure 5). Compared to the bulk soil, the clustering in the rhizoplane for ERGO and mycorrhizal treatments (ME) was significantly more compared to other treatments and sampling zones (Figure 5). In contrast, the PCoA analysis for fungal communities across the sampling zones and treatments showed a non-specific effect of the treatments and the location of microbiome sampling (Figure 6). There was no significant effect in the distribution and clustering of fungal communities across ERGO and mycorrhizal treatments (Figure 6).

### AMF and ERGO amendments on indicator species

Indicator Species Analysis (ISA) was conducted, and the results were visualized using multi-panel bubble plots and corresponding summary tables (See SUPPLEMENTARY ONLINE Figure 2-3; and SUPPLEMENTARY ONLINE Tables 1-2). Based on significant indicator values (IndVal), microbial taxa (both bacteria and fungi) were differentially enriched in plants inoculated with the ERGO/Mycorrhizal or both. The 16S indicator species plot highlights bacterial species whose abundance is significantly influenced by treatment and sampling zone, illustrating distinct microbiome shifts. The plot and table generated from the indicator species analysis highlight taxa with the highest indicator values, representing species strongly associated with specific treatments and sample types. In the 16S dataset, *Planctomycete A-2* exhibited notably high indicator values and were relatively abundant under the ME treatment across all sample types, indicating a strong and consistent association (See SUPPLEMENTARY ONLINE Figure 3-4; and SUPPLEMENTARY ONLINE Tables 1-2). Similarly, *Thioalkalivibrio* demonstrated high indicator strength in the ERGO treatment, although its relative abundance remained low across sample types. On the contrary, *Holophaga sps.* with the treatment ERGO was shown to be relatively more abundant but its indicator value was not a significant indicator for this treatment, suggesting it may be a rare but treatment-specific taxon. *Chlorobi* bacteria were exclusively detected in the ME treatment across root, rhizosphere, and bulk soil, marked by strong indicator values and consistently low abundance, underscoring their specificity to this treatment. *Flavobacterium caeni* displayed a relative abundance of approximately 0.04%, which was moderate under the M treatment in root samples but lower in ME-treated rhizosphere and soil samples. *Acidobacteria bacterium* showed moderate relative abundance across E, M, and ME treatments in all sample types. Among all sample types, the rhizosphere exhibited the highest species diversity and number of significant indicator taxa, followed by root and bulk soil, reflecting strong microbial community differentiation across plant-associated niches.

In contrast, the ITS-based indicator species analysis revealed a smaller number of significantly associated fungal taxa, *Metarhizium anisopliae* had high indicator values despite low relative abundance, particularly under the M treatment, indicating strong specificity to this treatment. *Trichomonascus valeenenianus* was moderately abundant and exhibited moderate indicator values in the E treatment, suggesting a consistent but less pronounced association. The *GS05* fungal group was more abundant under M treatment but showed varying degrees of association strength across samples. Notably, *Candida subhashi* was detected with relatively high abundance only in the ME treatment within soil samples, accompanied by moderately high indicator values, indicating its strong treatment and niche-specific presence. Interestingly, *Paecilomyces zollerniae* was present in both root and rhizosphere under the E treatment, with higher indicator values in the rhizosphere but relatively greater abundance in both sample types, suggesting niche-dependent indicator strength. (See SUPPLEMENTARY ONLINE Figure 2-3; and SUPPLEMENTARY ONLINE Tables 1-2)

### Influence on microbial co-occurrence network in rhizosphere microbiome

Correlation network analyses were performed separately for the M (AMF only) and ME (AMF + ERGO) treatments (Figure 7 and 8), to indentify bacterial associations driven by the presence of introduced AMF species. Treatment E was not included, since it lacks AMF addition. Each network was constructed using data combined from all sample types (root, rhizosphere, and soil), with grouping based solely on the treatment. The construction of the bacterial networks revealed that inoculation with mycorrhizae in wheat rhizosphere affected the complexity of the network (Figure 7). The number of edges in the network of plants inoculated with the AMF (M) was reduced compared with that in the network plants inoculated with both ERGO and AMF (ME) (Figure 7 and 8). A decrease in the number of nodes was also observed in inoculated plants, especially when the plants were inoculated with just AMF compared to a dual inoculum of AMF + ERGO (Figure 7 and 8). Moreover, a greater number of nodes, modularity, and number of communities were observed when plants were co-inoculated with AMF + ERGO (ME) (Figure 7 and 8; SUPPLEMENTARY ONLINE Tables 1-2). Interestingly, AMF only inoculation also created a more of negative correlation with few bacterial taxa (Figure 7). In contrast, compared with AMF treatments, AMF + ERGO (ME) inoculation resulted in a greater total number of edges, including negative and positive edges (Figure 8). Thus, compared to the AMF only treatment, more co-occurrences (both positive and negative) were observed with AMF + ERGO (ME) treatments. The correlation network analysis for the M treatment, filtered at FDR < 0.05, revealed robust and statistically significant associations between *Claroideoglomus etunicatum* and multiple bacterial genera. Notably, *C. etunicatum* exhibited strong positive correlations with *Aeromicrobium*, LWQ8, *Luteitalea*, *Aestuariimicrobium*, and *Cytophaga*, with no significant negative associations observed. Figure 7B illustrates the correlation network under the same treatment based on a less stringent unadjusted p-value < 0.05. This network uncovered additional, weaker associations, including negative correlations—highlighted by blue edges—between *Rhizophagus irregularis* and *Funneliformis mosseae* with certain bacterial taxa. *Verrucomicrobium* was negatively associated with both *R. irregularis* and *F. mosseae*, while *C. etunicatum* remained free of significant negative correlations.

**Figure 7:**
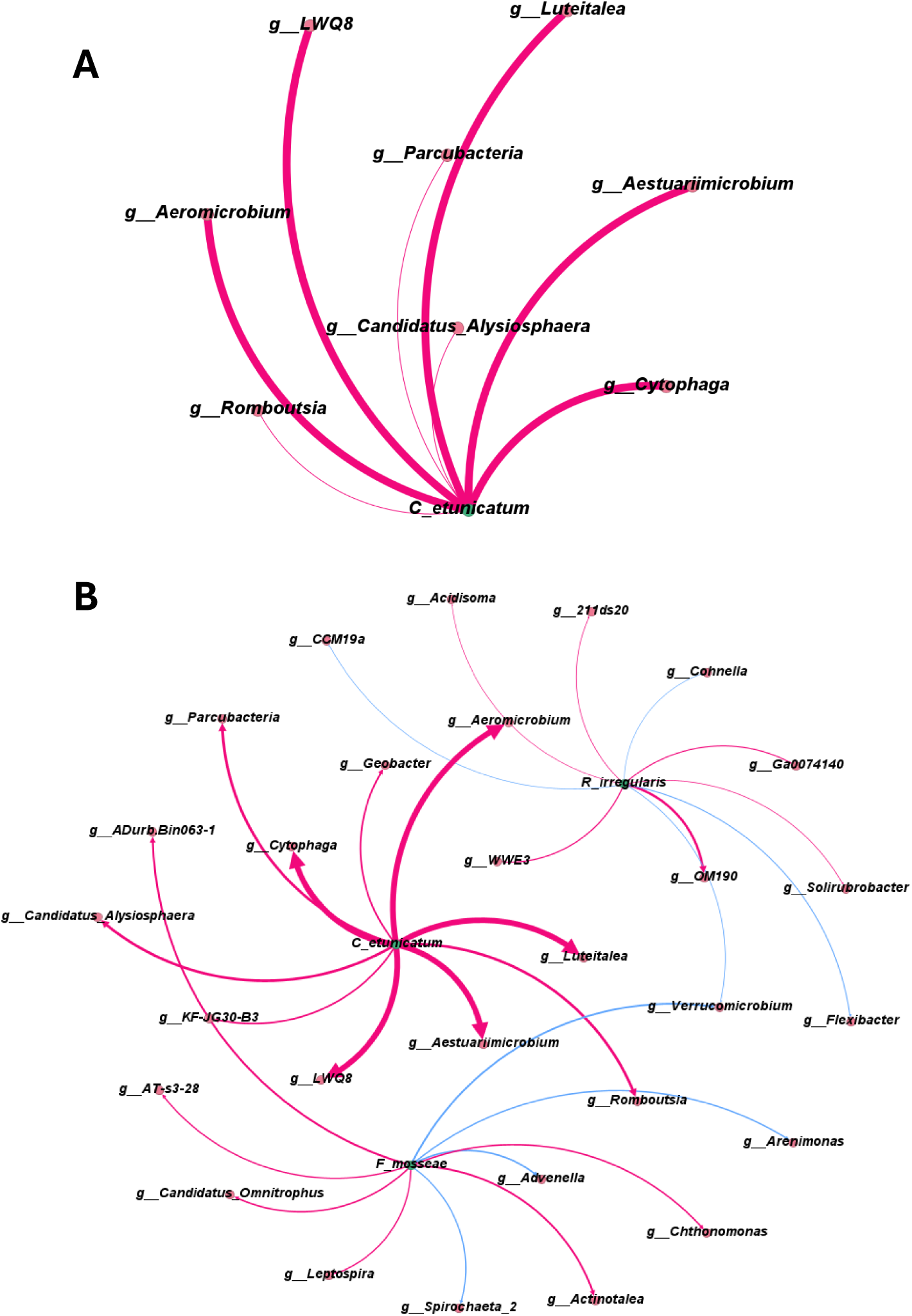
A Correlation network representing significant associations between AMF species and bacterial taxa in the samples treated with lone mycorrhizae (M), based on false discovery rate (FDR) P< 0.05. B Correlation network for the same treatment, based on unadjusted p-value < 0.05. Edge width indicates the strength of the correlation. Magenta edges represent positive correlations, while blue edges indicate negative correlations between bacterial taxa and AMF species. Blue nodes represent AMF species, and pink nodes represent bacterial taxa significantly associated with AMF species.

**Figure 8:**
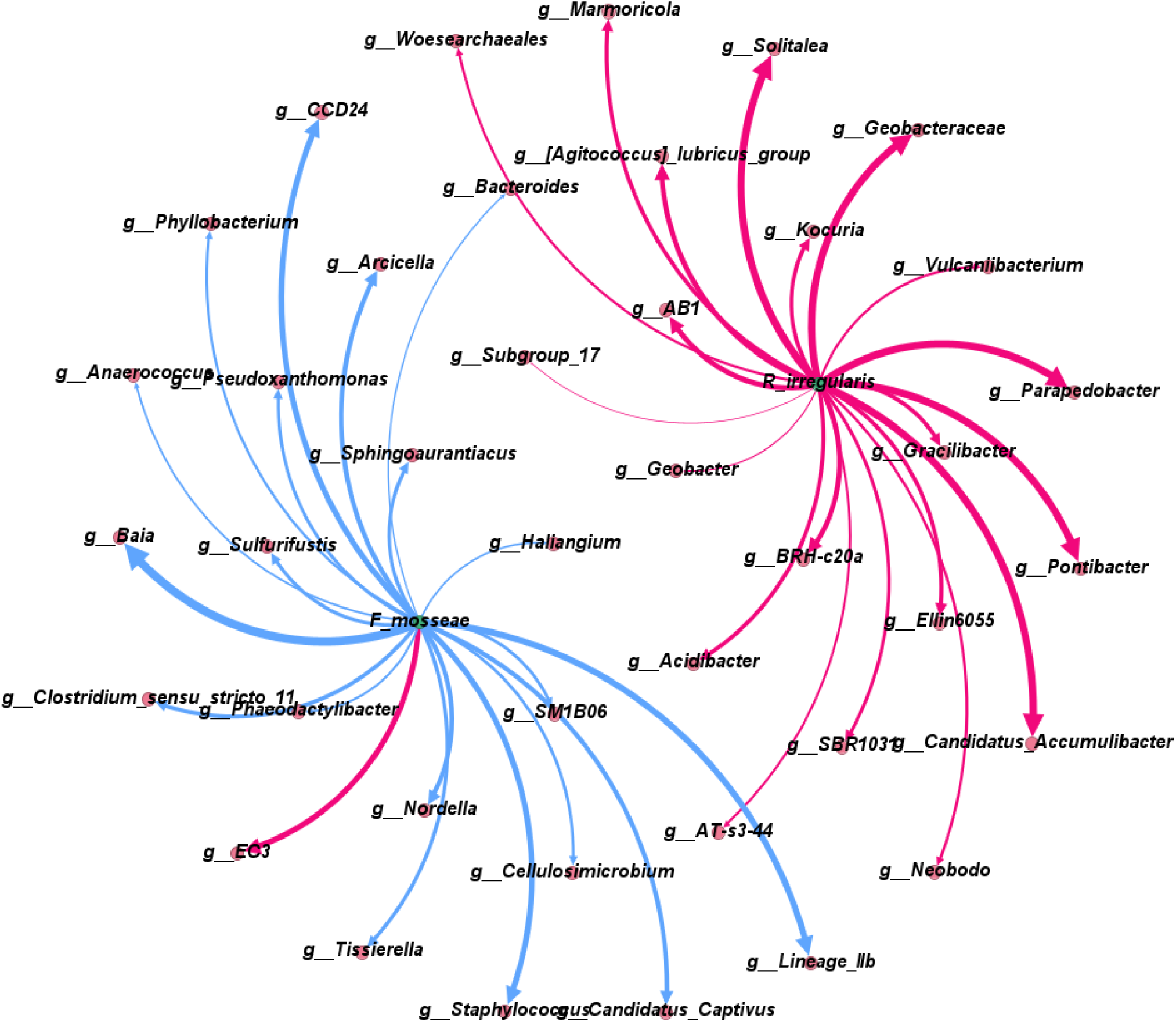
Correlation networks illustrating associations between AMF species and bacterial taxa in the AF + ERGO (ME) treatment group, based on unadjusted p-value < 0.05. Edge width corresponds to the strength of the correlation. Magenta edges indicate positive correlations, while blue edges indicate negative correlations. Green nodes represent AMF species, and pink nodes represent bacterial taxa significantly correlated with the respective AMF species.

Under the ME treatment, the correlation network indicated a shift in interaction patterns (Figure 8). *R. irregularis* showed strong positive correlations with several bacterial genera, whereas *F. mosseae* demonstrated primarily negative associations, with a single exception—*EC3*—exhibiting a positive correlation. No bacterial taxa were found to be co-associated with both AMF species in the ME treatment. These results highlight the distinct influence of AMF species on bacterial communities, with *C. etunicatum* emerging as a key contributor to shaping bacterial composition under the M treatment.

## Discussion

In nature, the minimal ERGO levels found in some plants suggest that arbuscular mycorrhizal fungi (AMF) may act as both a reservoir of ERGO biosynthesis and also a driver to facilitate ERGO mobilization in the plants. It is speculated that ERGO may either be directly synthesized by mycorrhizae or absorbed from the soil. Recent studies have shown that mycorrhizae, particularly AMFs, play a role in increasing ERGO content in grain, but it is currently unknown if that is due to uptake from the soil or biosynthesis by AMF [8]. These experiments, conducted under natural soil conditions, showed that the association of AMF with multiple crops led to the uptake of ERGO at the grain level [8]. It is argued that while mammalian systems do not synthesize ERGO, they can absorb it through known transporters [35]. It has recently been shown that invertebrate model systems, such as *Caenorhabditis elegans,* can take up ERGO, leading to increased lifespan and overall health in the worms [36]. While the plant uptake of ERGO remains unknown, the abundance of AMFs in soil suggests that plants, in association with soil fungi, may facilitate the uptake of ERGO.

Currently, the uptake and storage of ERGO in plants are not fully understood. Therefore, the inoculation of pure ERGO into soil with AMF may encounter various challenges when interacting with the natural soil microbiome due to the diverse microbial community and the complexity of the relationships present around the roots [37]. It is now recognized that plants actively shape the rhizosphere microbiome, and the regulation of microbiome diversity around the roots also influences the overall plant developmental phenotype. [38]

Wheat plants inoculated with ERGO exhibited better growth traits compared to the untreated control, suggesting the beneficial role of ERGO. This growth-promoting effect of ERGO parallels similar observations in both invertebrate and mammalian model systems, where ERGO has been shown to promote growth, lifespan, and other health benefits [39]. The observation of growth benefits from direct ERGO supplementation (only) also indicated that plants could take up ERGO in the absence of AMFs, suggesting the involvement of a receptor or transporter for the temporal uptake of ERGO. Interestingly, direct ERGO supplementation in the wheat rhizosphere led to temporal uptake of ERGO in the roots, shoots, and grains. The ERGO concentration in roots was higher in both ERGO and AMF treatments compared to the AMF + ERGO treatments, suggesting that plant association with AMFs may facilitate ERGO uptake, as observed in other studies [8]. Similarly, the direct uptake of ERGO when pure ERGO was supplied to the roots suggests that plants can take up ERGO both with and without AMFs, indicating two distinct mechanisms of ERGO uptake. In contrast, grain fortification with ERGO was higher in the AMF + ERGO treatment compared to individual AMF and ERGO treatments, suggesting the existence of distinct transport mechanisms for ERGO grain fortification, separate from those involved in shoot/root uptake.

It is shown that amino acid content in the rhizosphere changes based on the plants’ ability to secrete amino acids or microbial transformation of amino acids in soil [40]. ERGO is an unusual amino acid known to be biosynthesized by fungal species and few actinobacteria [17]. ERGO is a dietary thione/thiol and it improves lifespan and muscle density in both invertebrate and mammalian model systems [36]. It is reasoned that ERGO improves traits in both invertebrates and mammalian model systems by augmenting the microbiome [41]. However, the role of ERGO, either synthesized by soil fungi or by direct supplementation on soil microbiome, is not investigated. It is shown that the microbiome diversity in the different rhizosphere zones plays a critical role in plant performance [38]. The bacterial microbiome (both diversity and structure) was more influenced by ERGO and AMF treatments in the rhizosphere compared to the root plane, suggesting the role of ERGO in modulating bacterial community in the rhizosphere. Concurrently, bacteria and fungi were enriched in the presence of both ERGO and AMF treatments.

The microbial responses observed under ergothioneine (ERGO) treatment reveal diverse ecological and metabolic adaptations across bacterial and fungal taxa. *Thioalkalivibrio* species exhibited a high indicator value under ERGO treatment. This aligns with earlier findings [42], wherein EgtC function was characterized in the involment of ERGO biosynthesis. EgtC-like homologs identified in *Thioalkalivibrio,* a Gram-negatve bacterial genus, suggest a potential role in ERGO-related metabolism, even in the absence of canonical ERGO biosynthetic genes. In addition, our results showed increased abunance of *Holophaga* (a member of the phylum Acidobacteria) under ERGO treatment. A previous study indicate that [43], *Holophaga foetida* utilizes methoxylated aromatic compounds in lignin of plant cell walls through transmethylation processes, producing volatile sulfur compounds. This metabolic specialization may confer an advantage in ERGO-enriched environments, possibly due to structural similarities between ERGO or its derivatives and the substrates preferred by *Holophaga*. Further, a Chlorobi-affiliated bacterium was predominantly present in the root compartment under ERGO treatment [44]. The study identified an oxygen-independent ERGO biosynthesis pathway in *Chlorobium limicola*, mediated by the enzyme EanB [44]. This pathway enables ERGO production under anaerobic conditions, characteristic of the root microenvironment. The high indicator value of this *Chlorobi* bacterium suggests a strong association with ERGO-enriched conditions, potentially reflecting its metabolic adaptation to utilize or produce ERGO in anoxic niches.

*Paecilomyces zollerniae* emerged as an indicator species under ERGO treatment in ITS Indicator species analysis. While not directly studied for ERGO biosynthesis, other species within the *Paecilomyces* genus, such as *Paecilomyces tenuipes*, are known producers of ERGO in fruiting bodies[45]ERGO levels in *Paecilomyces tenuipes* increase significantly when cultures are supplemented with methionine, a known precursor in ERGO biosynthesis. Although *P. zollerniae* and *P. tenuipes* differ taxonomically, they share potential for similar metabolic capabilities. Enriched *P. zollerniae* under ERGO treatment may reflect either a tolerance to or utilization of ERGO, highlighting possible unexplored biosynthetic potential within this genus. Collectively, these observations underscore the multifaceted roles of ERGO in microbial ecology, influencing community composition and metabolic functions across diverse taxa (SUPPLEMENTARY FIGURES 2-3; SUPPLEMENTARY TABLE 1-2). These findings demonstrate that ERGO, both independently and in combination with AMF, significantly influenced microbiome composition and diversity.

In this study, AMF treatment also significantly influences microbial composition. *Claroideoglomus etunicatum*, one of the applied arbuscular mycorrhizal fungi (AMF) species, has been shown to enhance ERGO uptake in plants [8]. The robust positive correlations observed between *C. etunicatum* (see Figure 7 A) and certain bacterial species under AMF lone treatment suggest synergistic interactions that may enhance ERGO availability and uptake. The genus Staphylococcus, known to harbor the EgtU transporter for ergothioneine uptake [46], was negatively correlated with the presence of *Funneliformis mosseae* one of the added AMF species in the ERGO and AMF treatment (Figure 8), suggesting that the AMF inoculation may alter microbial community dynamics in a way that suppresses EGT-utilizing taxa. One possible explanation is that AMF improves rhizosphere health and reduces oxidative stress, thereby diminishing the ecological advantage of EGT uptake. Alternatively, AMF may promote microbial competitors or alter the availability of sulfur-based metabolites, indirectly limiting the niche suitable for Staphylococcus. These findings highlight a potential antagonistic interaction between AMF and opportunistic EGT-utilizing bacteria.

## Conclusion

This study demonstrates that ergothioneine (ERGO) and arbuscular mycorrhizal fungi (AMF), individually and in combination, enhance wheat growth and significantly influence rhizosphere microbial communities. ERGO uptake occurred across plant tissues, with dynamics modulated by AMF presence, suggesting multiple transport pathways. Co-application of ERGO and AMF not only boosted biomass and grain yield but also increased bacterial diversity and reshaped microbial community structures, particularly through synergistic effects. Indicator species and microbial network analyses further revealed treatment-specific associations, indicating that AMF may help mediate beneficial microbial interactions while modulating EGT-utilizing taxa. These findings highlight the potential of ERGO and AMF as bioinoculants to improve crop productivity and soil microbiome function, offering a promising strategy for sustainable agriculture.

### Materials and data availability

All the analyzed datasets including 16s, ITS ASVs, metadata and R scripts regarding this research are included in this article as its supplementary information and are deposited in a publicly available in the following github repository. https://github.com/srisnandini/AMF-ERGO-.

## Supporting information

SOM Figures 1-3

SOM Table 1

SOM Table 2

## Acknowledgements

The work was conducted with the grant received by authors from USDA FFAR.

## Legends to Supplementary online Figures

**Figure SOM 1:** Representative images from the treatment groups and control on days 40, 50, and 60 of harvest. Plants were inoculated with either ergothioneine (ug per pot) or mycorrhizae (x no spores) or were inoculated with both the treatments. Control samples are not treated with either mycorrhizal species or ergothioneine. The other treatment depicts: ‘+M+E’ treatment in represents wheat plants inoculated with both mycorrhizal species and ergothioneine. ‘E’ treatment represents wheat plants inoculated with ergothioneine. ‘M’ treatment in represents wheat plants inoculated with mycorrhizal species.

**Figure SOM 2:** The bubble plot highlights: Indicator species analysis of the top 20 16S bacterial community across three sample types—Root, Rhizosphere, and Soil—under four treatment conditions (Control, E, M, ME). The species with significant indicator values (IndVal), arranged in descending order along the y-axis. Each bubble denotes a species’ association with a treatment group. Bubble size represents the average relative abundance (%), while bubble color reflects statistical significance, with darker shades indicating lower p-values. Rhizosphere samples exhibit greater species diversity and higher indicator values than root and soil. Only taxa with p-values < 0.05 are shown, emphasizing species most responsive to treatment effects.

**Figure SOM 3: T**he bubble plot highlights: Indicator species analysis of the top 20 ITS bacterial community across three sample types—Root, Rhizosphere, and Soil—under four treatment conditions (Control, E, M, ME). The species with significant indicator values (IndVal), arranged in descending order along the y-axis. Each bubble denotes a species’ association with a treatment group. Bubble size represents the average relative abundance (%), while bubble color reflects statistical significance, with darker shades indicating lower p-values. Rhizosphere samples exhibit greater species diversity and higher indicator values than root and soil. Only taxa with p-values < 0.05 are shown, emphasizing species most responsive to treatment effects.

## References

1. Styczen ME, Abrahamsen P, Hansen S, Knudsen L. Model analysis of the significant drop in protein content in Danish grain crops from 1990-2015. Eur. J. Agron. 2020; 118:126068.

2. Simmonds NW. The relation between yield and protein in cereal grain. J. Sci. Food Agric.. 1995;67(3):309–15.

3. Dong J, Gruda N, Lam SK, Li X, Duan Z. Effects of elevated CO2 on nutritional quality of vegetables: a review. Front. Plant Sci. 2018; 9:924.

4. Malila Y, Owolabi IO, Chotanaphuti T, Sakdibhornssup N, Elliott CT, Visessanguan W, Karoonuthaisiri N, Petchkongkaew A. Current challenges of alternative proteins as future foods. npj Sci Food. 2024;8(1):53.

5. Berg G, Rybakova D, Fischer D, Cernava T, Vergès MC, Charles T, Chen X, Cocolin L, Eversole K, Corral GH, Kazou M. Microbiome definition re-visited: old concepts and new challenges. Microbiome. 2020; 8:1–22.

6. Qi S, Wang J, Wan L, Dai Z, da Silva Matos DM, Du D, Egan S, Bonser SP, Thomas T, Moles AT. Arbuscular mycorrhizal fungi contribute to phosphorous uptake and allocation strategies of Solidago canadensis in a phosphorous-deficient environment. Front. Plant Sci. 2022; 13:831654.

7. Whiteside MD, Garcia MO, Treseder KK. Amino Acid Uptake in Arbuscular Mycorrhizal Plants. PLoS ONE. 2012; 7(10): e47643.

8. Carrara JE, Lehotay SJ, Lightfield AR, Sun D, Richie Jr JP, Smith AH, Heller WP. Linking soil health to human health: Arbuscular mycorrhizae play a key role in plant uptake of the antioxidant ergothioneine from soils. PPP. 2023;5(3):449–58.

9. Borodina I, Kenny LC, McCarthy CM, Paramasivan K, Pretorius E, Roberts TJ, van der Hoek SA, Kell DB. The biology of ergothioneine, an antioxidant nutraceutical. Nutr Res Rev. 2020;33(2):190–217.

10. Beelman RB, Richie Jr JP, Phillips AT, Kalaras MD, Sun D, Duiker SW. Soil disturbance impact on crop ergothioneine content connects soil and human health. Agron. 2021;11(11):2278.

11. Halliwell B, Cheah IK, Tang RM. Ergothioneine–a diet-derived antioxidant with therapeutic potential. FEBS Lett. 2018;592(20):3357–66.

12. Fu TT, Shen L. Ergothioneine as a natural antioxidant against oxidative stress-related diseases. Front. Pharmacol. 2022; 13:850813.

13. Melville DB, Eich S. The occurrence of ergothioneine in plant material. J. Biol. Chem, 1956;218(2), 647–651.

14. Bello MH, Barrera-Perez V, Morin D, Epstein L. The Neurospora crassa mutant NcΔEgt-1 identifies an ergothioneine biosynthetic gene and demonstrates that ergothioneine enhances conidial survival and protects against peroxide toxicity during conidial germination. Fungal Genet Biol. 2012;49(2):160–72.

15. Hu W, Song H, Sae Her A, Bak DW, Naowarojna N, Elliott SJ, Qin L, Chen X, Liu P. Bioinformatic and biochemical characterizations of C–S bond formation and cleavage enzymes in the fungus Neurospora crassa ergothioneine biosynthetic pathway. Org. Lett. 2014;16(20):5382–5.

16. Seebeck FP. In vitro reconstitution of mycobacterial ergothioneine biosynthesis.J Am Chem Soc. 2010;132(19):6632–3.

17. Cumming BM, Chinta KC, Reddy VP, Steyn AJ. Role of ergothioneine in microbial physiology and pathogenesis. Antioxid. Redox Signal. 2018;28(6):431–44.

18. Paul BD. Ergothioneine: a stress vitamin with antiaging, vascular, and neuroprotective roles. Antioxid. Redox Signal. 2022;36(16-18):1306–17

19. Gründemann D, Harlfinger S, Golz S, Geerts A, Lazar A, Berkels R, Jung N, Rubbert A, Schömig E. Discovery of the ergothioneine transporter.Proc Natl Acad Sci.. 2005;102(14):5256–61.

20. Parada AE, Needham DM, Fuhrman JA. Every base matter: assessing small subunit rRNA primers for marine microbiomes with mock communities, time series and global field samples. Environ. Microbiol. 2016;18(5):1403–14.

21. Apprill A, McNally S, Parsons R, Weber L. Minor revision to V4 region SSU rRNA 806R gene primer greatly increases detection of SAR11 bacterioplankton.Aquat. Microb. Ecol. 2015;75(2):129–37.

22. Gilbert JA, Meyer F, Jansson J, Gordon J, Pace N, Tiedje J, Ley R, Fierer N, Field D, Kyrpides N, Glöckner FO. The earth microbiome project: meeting report of the “1 st EMP meeting on sample selection and acquisition” at Argonne National Laboratory October 6 th 2010. SIGS. 2010; 3:249–53.

23. White TJ, Bruns T, Lee SJ, Taylor J. Amplification and direct sequencing of fungal ribosomal RNA genes for phylogenetics. PCR protocols: a guide to methods and applications. RG. 1990;18(1):315–22.

24. Bolyen E, Rideout JR, Dillon MR, Bokulich NA, Abnet CC, Al-Ghalith GA, Alexander H, Alm EJ, Arumugam M, Asnicar F, Bai Y. Reproducible, interactive, scalable and extensible microbiome data science using QIIME 2. Nat. Biotechnol. 2019;37(8):852–7.

25. Quast C, Pruesse E, Yilmaz P, Gerken J, Schweer T, Yarza P, Peplies J, Glöckner FO. The SILVA ribosomal RNA gene database project: improved data processing and web-based tools. Nucleic Acids Res. 2012;41(D1):D590–6.

26. Ssekagiri AT, Sloan W, Ijaz UZ. microbiomeSeq: an R package for analysis of microbial communities in an environmental context. RG. 2017;(Vol. 10).

27. Shannon CE. A mathematical theory of communication. The Bell Syst. Tech J. 1948 Jul;27(3):379–423

28. McMurdie PJ, Holmes S. phyloseq: an R package for reproducible interactive analysis and graphics of microbiome census data. PloS one. 2013;8(4):e61217.

29. Anderson MJ. Permutational multivariate analysis of variance (PERMANOVA). Wiley statsref: statistics reference online. RG. 2014;1–5.

30. Severns PM, Sykes EM. Indicator species analysis: A useful tool for plant disease studies. Phytopathol. 2020;110(12):1860–2.

31. Demircan T, Ovezmyradov G, Yıldırım B, Keskin İ, İlhan AE, Fesçioğlu EC, Öztürk G, Yıldırım S. Experimentally induced metamorphosis in highly regenerative axolotl (ambystoma mexicanum) under constant diet restructures microbiota. Sci. Rep. 2018;8(1):10974.

32. Harrell Jr FE, Harrell Jr MF. Package ‘hmisc’. CRAN2018. 2019; 2019:235–6.

33. Bastian M, Heymann S, Jacomy M. Gephi: an open-source software for exploring and manipulating networks. InProceedings of the international AAAI conference on web and social media 2009;(Vol. 3, No. 1, pp. 361–362).

34. Gajdoš P, Ježowicz T, Uher V, Dohnálek P. A parallel Fruchterman–Reingold algorithm optimized for fast visualization of large graphs and swarms of data. Swarm and evolutionary computation. RG. 2016; 26:56–63.

35. Shimizu T, Masuo Y, Takahashi S, Nakamichi N, Kato Y. Organic cation transporter Octn1-mediated uptake of food-derived antioxidant ergothioneine into infiltrating macrophages during intestinal inflammation in mice. DMPK. 2015;30(3):231–9.

36. Petrovic D, Slade L, Paikopoulos Y, D’Andrea D, Savic N, Stancic A, Miljkovic JL, Vignane T, Drekolia MK, Mladenovic D, Sutulovic N. Ergothioneine improves healthspan of aged animals by enhancing cGPDH activity through CSE-dependent persulfidation. Cell Metab. 2025;37(2):542–56.

37. Li J, Zhou L, Chen G, Yao M, Liu Z, Li X, Yang X, Yang Y, Cai D, Tuerxun Z, Li B. Arbuscular mycorrhizal fungi enhance drought resistance and alter microbial communities in maize rhizosphere soil. Environ. Technol. Innov. 2025; 37:103947.

38. Pantigoso HA, Newberger D, Vivanco JM. The rhizosphere microbiome: Plant–microbial interactions for resource acquisition. J. Appl. Microbiol. 2022;133(5):2864–76.

39. Katsube M, Ishimoto T, Fukushima Y, Kagami A, Shuto T, Kato Y. Ergothioneine promotes longevity and healthy aging in male mice. GeroScience. 2024;46(4):3889–909.

40. Moe LA. Amino acids in the rhizosphere: from plants to microbes. Am. J. Bot. 2013;100(9):1692–705.

41. Matsuda Y, Ozawa N, Shinozaki T, Wakabayashi KI, Suzuki K, Kawano Y, Ohtsu I, Tatebayashi Y. Ergothioneine, a metabolite of the gut bacterium Lactobacillus reuteri, protects against stress-induced sleep disturbances. Transl. Psychiatry. 2020;10(1):170.

42. Vit A, Mashabela GT, Blankenfeldt W, Seebeck FP. Structure of the ergothioneine-biosynthesis amidohydrolase EgtC. ChemBioChem. 2015;16(10):1490–6.

43. Liesack W, Bak F, Kreft JU, Stackebrandt E. Holophaga foetida gen. nov., sp. nov., a new, homoacetogenic bacterium degrading methoxylated aromatic compounds. Arch. Microbiol. 1994; 162:85–90.

44. Burn R, Misson L, Meury M, Seebeck FP. Anaerobic Origin of Ergothioneine. Angew Chem Int Ed Engl. 2017;56(41):12508–12511

45. Lee WY, Park EJ, Ahn JK, Ka KH. Ergothioneine contents in fruiting bodies and their enhancement in mycelial cultures by the addition of methionine. Microbiol. 2009;37(1):43–7.

46. Zhang Y, Gonzalez-Gutierrez G, Legg KA, Walsh BJ, Pis Diez CM, Edmonds KA, Giedroc DP. Discovery and structure of a widespread bacterial ABC transporter specific for ergothioneine. Nat. Commun. 2022;13(1):7586.

