## Supplementary material for "Soil to human health continuum: Exploring ergothioneine and mycorrhizal fungi in shaping the wheat microbiome": SOM Figures 1-3

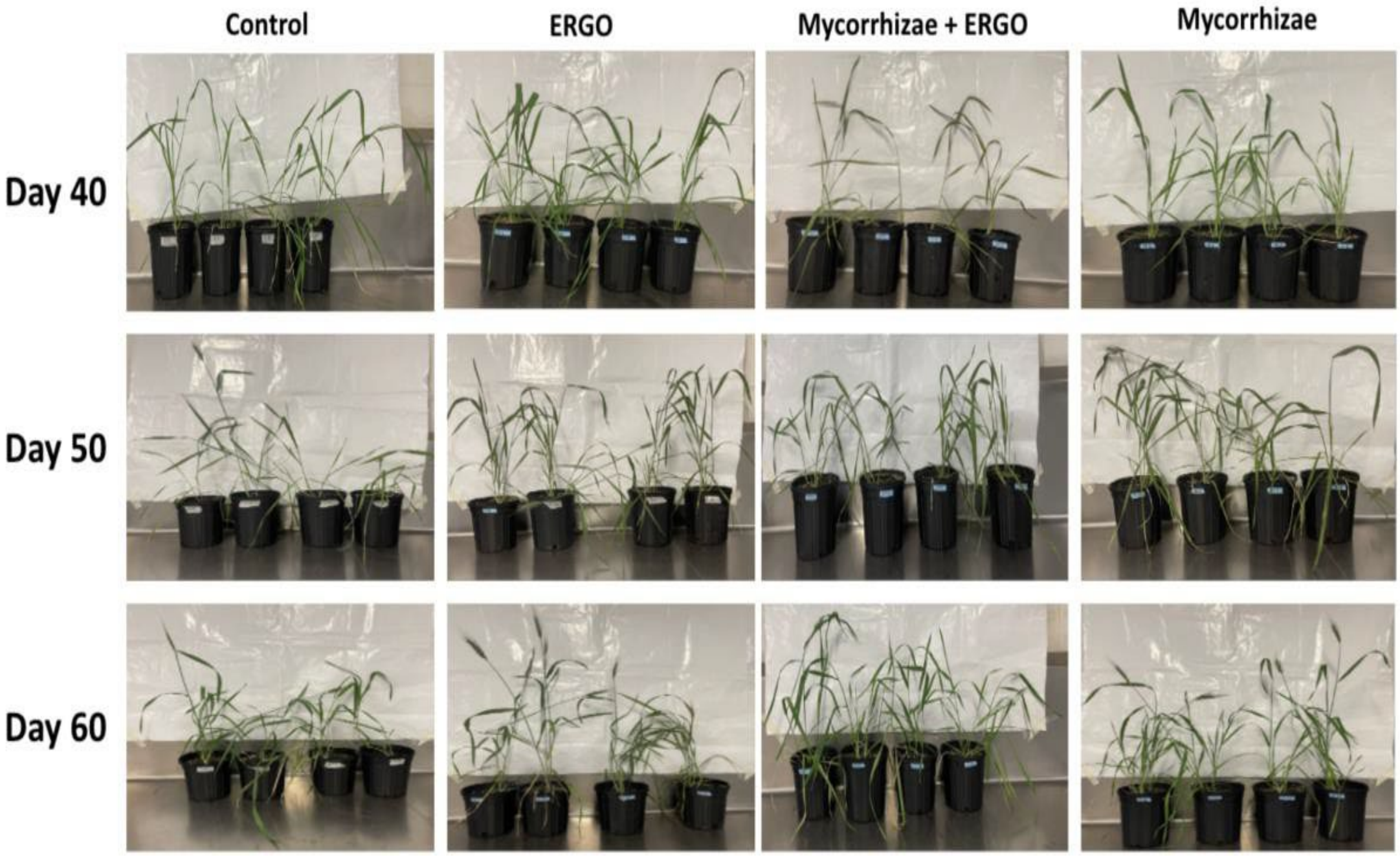


**Figure SOM 1:** Representative images from the treatment groups and control on days 40, 50, and 60 of harvest. Plants were inoculated with either ergothioneine (ug per pot) or mycorrhizae (x no spores) or were inoculated with both the treatments. Control samples are not treated with either mycorrhizal species or ergothioneine. Control samples are not treated with either mycorrhizal species or ergothioneine. The other treatment depicts: 'ME' treatment represents wheat plants inoculated with both mycorrhizal species and ergothioneine; 'E' treatment represents wheat plants inoculated with ergothioneine; 'M' treatment represents wheat plants inoculated with mycorrhizal species.

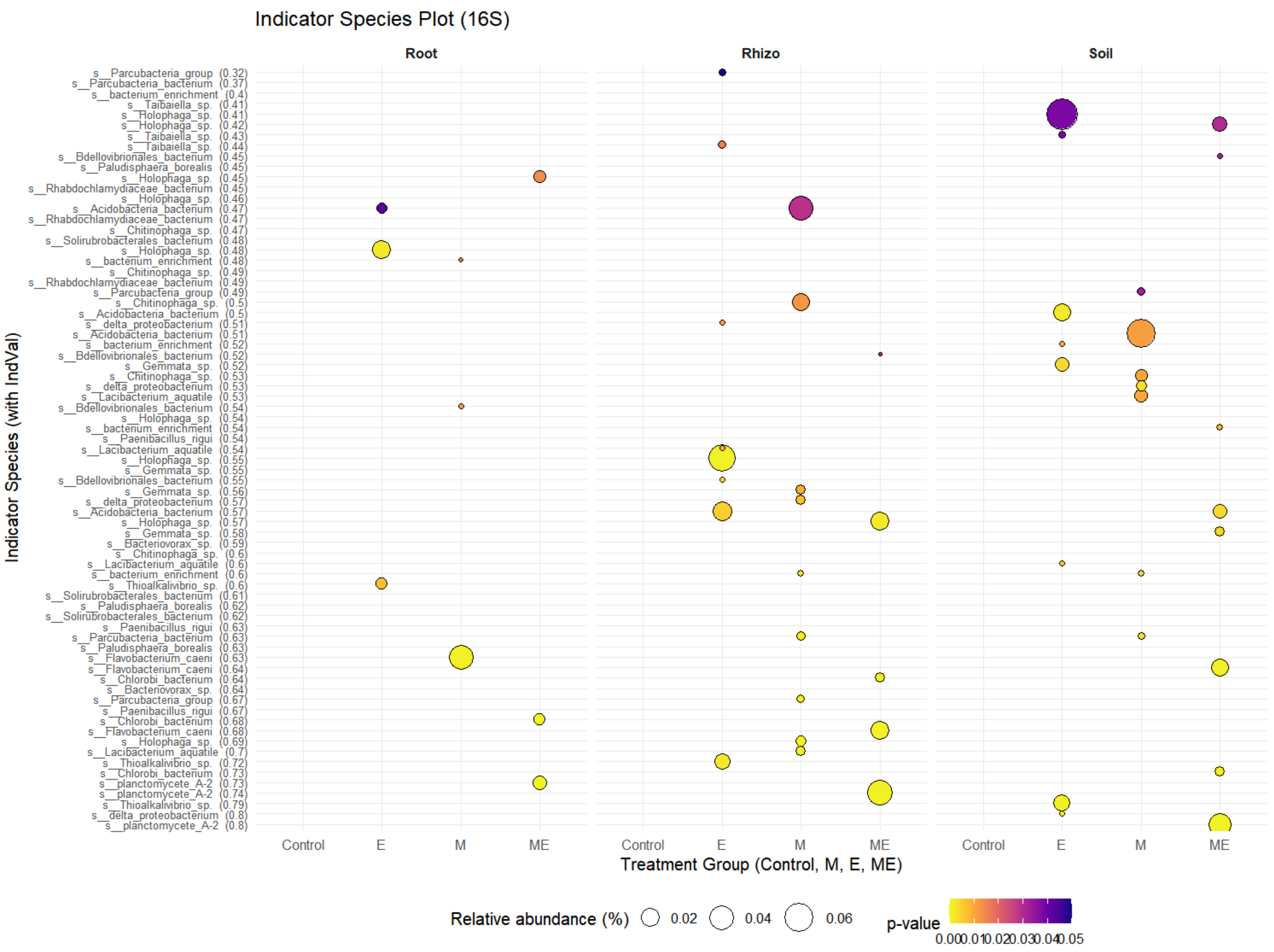


**Figure SOM 3:** Indicator species analysis to show the top 20 bacterial taxa across three sample types—Root, Rhizosphere, and Soil—under four treatment conditions (Control, E, M, ME). The species with significant indicator values (IndVal), arranged in descending order along the y-axis. Each bubble denotes a species' association with a treatment group. Bubble size represents the average relative abundance (%), while bubble color reflects statistical significance, with darker shades indicating lower p-values. Only taxa with p-values < 0.05 are shown.

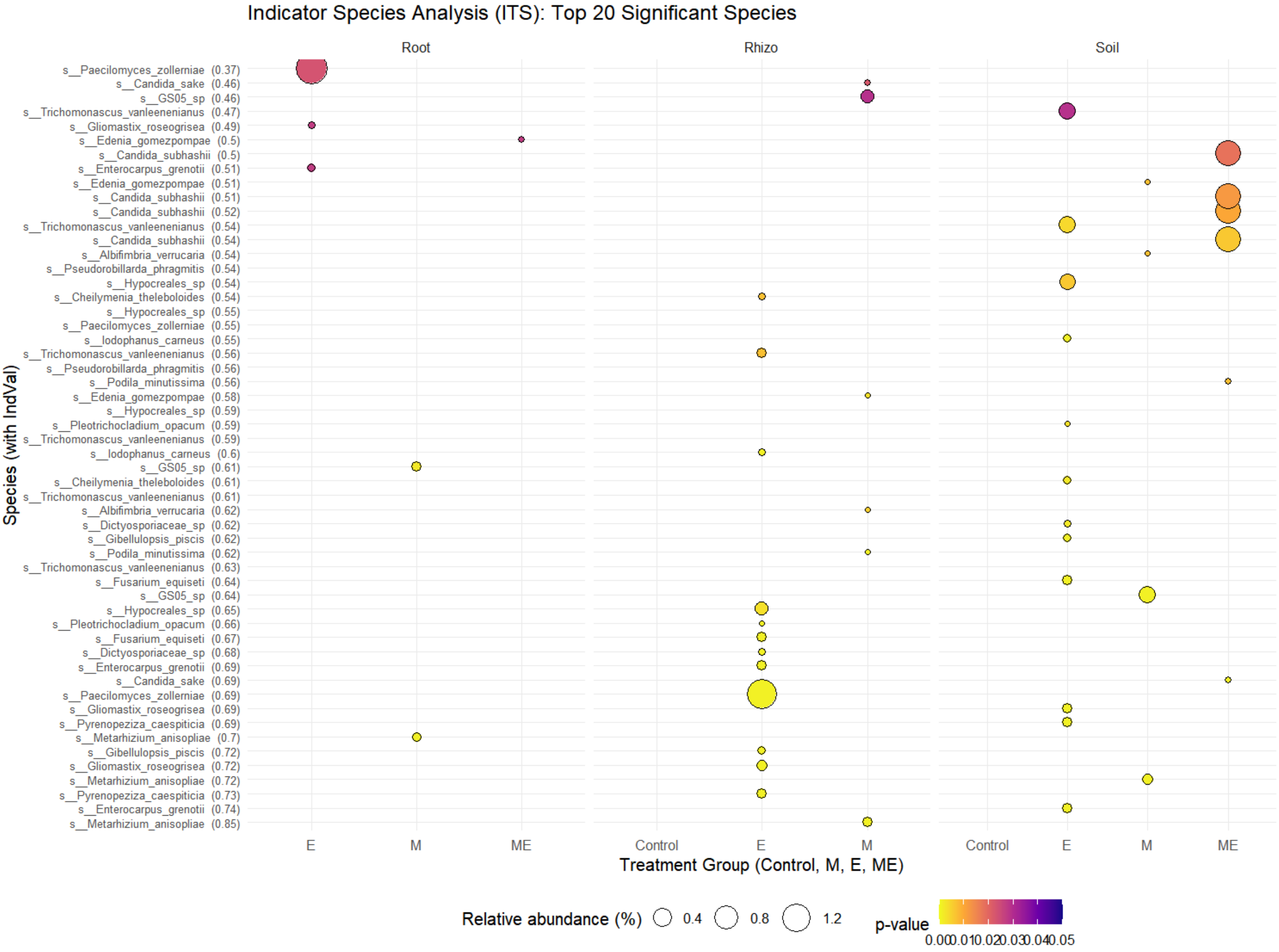


**Figure SOM 4:** Indicator species analysis to show the top 20 fungal taxa across three sample types—Root, Rhizosphere, and Soil—under four treatment conditions (Control, E, M, ME). The species with significant indicator values (IndVal), arranged in descending order along the y-axis. Each bubble denotes a species' association with a treatment group. Bubble size represents the average relative abundance (%), while bubble color reflects statistical significance, with darker shades indicating lower p-values. Only taxa with p-values < 0.05 are shown.
